# Reduced context updating but intact visual priors in autism

**DOI:** 10.1101/2021.10.21.464885

**Authors:** R. Randeniya, I. Vilares, J. B. Mattingley, M. I. Garrido

**Author notes:** Correspondence should be addressed to: Roshini Randeniya. **Disclosures** The authors have declared that there are no conflicts of interest in relation to the subject of this study.

## Abstract

A general consensus persists that sensory-perceptual differences in autism, such as hypersensitivities to light or sound, result from an overreliance on new (rather than prior) sensory observations. However, conflicting Bayesian accounts of autism remain unresolved as to whether such alterations are caused by more precise sensory observations (precise likelihood model) or by forming a less precise model of the sensory context (hypo-priors model). We used a decision-under-uncertainty paradigm that manipulated uncertainty in both likelihoods and priors. Contrary to model predictions we found no differences in reliance on likelihood in autistic group (AS) compared to neurotypicals (NT) and found no differences in subjective prior variance between groups. However, we found reduced context adjustment in the AS group compared to NT. Further, the AS group showed heightened variability in their relative weighting of sensory information (vs. prior) on a trial-by-trial basis. When participants were aligned on a continuum of autistic traits, we found no associations with likelihood reliance or prior variance but found an increased likelihood precision with autistic traits. These findings together provide empirical evidence for intact priors, precise likelihood, reduced context updating and heightened variability during sensory learning in autism.

## Introduction

Sensory processing alterations, which can affect one or more sensory modalities (Tavassoli et al, 2014b), are reported in around 90% of autistic adults (Crane et al, 2009). Autistic individuals can show improved performance in visual search tasks (Plaisted et al., 1998b; Joseph et al., 2009) and visual and auditory target detection (Mottron et al., 2000), compared to neurotypicals. However, sensory processing dysfunction (e.g. hypo- or hyper-sensitivities) can inhibit participation in activities such as learning and social interactions, which in turn impose lifelong challenges (Suarez, 2012). Due to the heterogeneous nature of perceptual function in autism, current diagnostic measures, such as the Autism Diagnostic Observation Schedule (ADOS), have significant limitations in characterizing the nature of perceptual experiences in people with a diagnosis of Autism Spectrum Disorder (ASD), and can only be applied for specific degrees of severity and age along the autism spectrum (Haker et al, 2016). Thus, better characterization of the mechanisms that give rise to autistic perception should help in understanding perceptual subtypes and could pave the way to personalized interventions.

“Bayesian brain” accounts for autism have offered explanations for perceptual differences in autism (Brock, 2012; Haker et al, 2016; Lawson et al, 2014; Pellicano & Burr, 2012; Van de Cruys et al, 2014). Simply stated, a Bayesian approach to sensory learning posits that during learning individuals form models, encoded as priors, by detecting patterns in the environment (Figure 1A). New incoming information (i.e., likelihood) is then matched against these priors. The internal model (prior) about the sensory environment is updated with new sensory information until model updating is no longer necessary in a stable environment (Penny, 2012). This framework has proven useful in explaining typical sensory learning and decision-making under ambiguity. According to Bayes theorem, posteriors (or perceptual decisions) will combine both prior and likelihood information but give more weight to whichever source has a higher precision (i.e., the lower variance). In autism, however, it is theorised that this process may be altered due to an imbalance of precision ascribed to sensory observations relative to prior beliefs (Lawson et al, 2014); (Note we use *precision* here in its statistical sense, which is the inverse of variance). From a Bayesian perspective, this likelihood over-reliance can be formalised mathematically as a shift in the posterior toward the sensory observation (likelihood) and away from the prior (belief). This shift can be attributed to different underlying causes. The “hypo-priors model” ((Pellicano & Burr, 2012); Figure 1B) suggests that the shift is caused by noisier or less precise (high variance) priors, equivalent to a weak model (or belief) of the environment. The “precise likelihood model” ((Brock, 2012); Figure 1C) argues instead that priors are intact, but there is an increase in the precision associated with sampling new information, such that sensory representations are more narrowly tuned. The conundrum is that given the relative difference in the precision of priors and likelihoods under the hypo-priors and precise likelihood models, the two accounts effectively give rise to the same posterior means.

**Figure 1:**
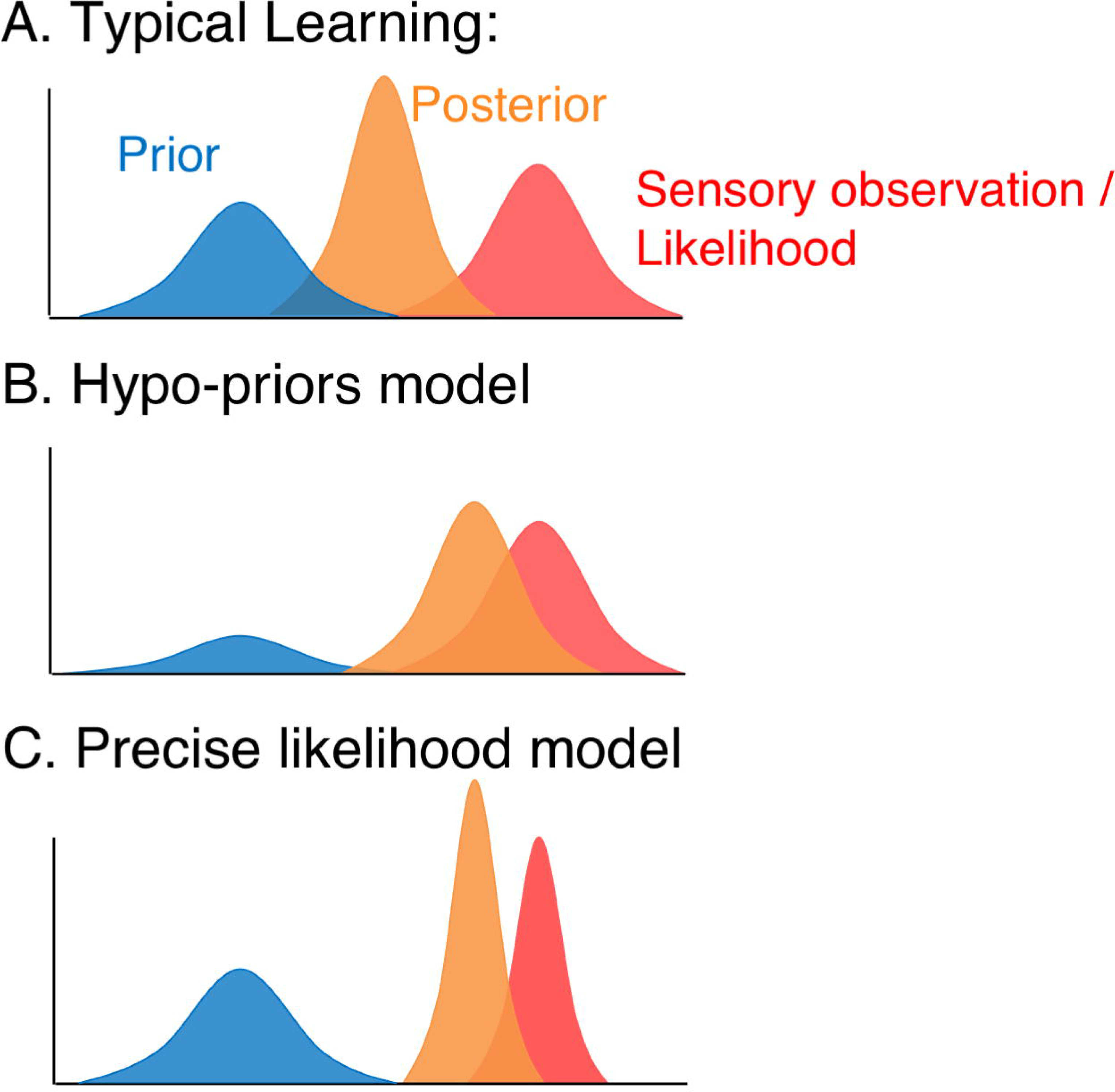
Bayesian models of atypical perception in Autism. A) Bayesian models postulate that in typical learning new sensory observations (likelihood; red) are integrated with the learned model of the world (Prior; blue), leading to a perception/decision (posterior; orange). B) Hypo-prior model for Autism (Pellicano & Burr, 2012): A weak model of the world (high variance in the prior) increases the reliance on new sensory observations (likelihood) that are relatively more precise than the prior (but as precise in autistic as in typical individuals) C) Precise Likelihood Model (Brock, 2012): Overly precise new sensory observations (likelihood has less variance than in typical individuals, and more precise than the prior) can increase the reliance on new sensory observations.

Amongst the criticisms of these Bayesian theories, it has been argued that while they can explain hypersensitivities and sensory overload, they do not account for hyposensitivity or other perceptual disruptions in autism such as weak global coherence (Teufel et al, 2013; Van de Cruys et al, 2014). Some studies investigating aberrant precision models in autism have found that children and adults on the autism spectrum are able to learn priors (Croydon et al, 2017; Pell et al, 2016), while other studies have shown that this process is altered (Skewes et al, 2015). Evidence for the precise likelihood model (Figure 1C) has been supported by a study showing that increasing autistic traits correlated with increasing precision in likelihood in a neurotypical sample (Karvelis et al., 2018), but not in prior representations. In contrast, another study which employed a signal detection approach provided evidence in support of the (weak) hypo-priors model (Skewes et al, 2015), again in a sample of neurotypical individuals. Further a study investigating central tendency in autistic children demonstrated poorer performance in autistic children than matched controls (Karaminis et al, 2016) indicating evidence for a hypo-priors model.In this study, we took a Bayesian approach to better understand how prior and likelihood information is utilized during visual sensory learning and perceptual decision-making in autism. Specifically, we investigated: 1) whether AS individuals rely more on sensory than on prior information compared with neurotypicals (NT), and 2) whether precision in prior and likelihood distributions differs between AS and NT groups. Further, given the utility of undertaking a hybrid of categorical and dimensional approaches to understanding autism (Abu-Akel et al, 2019; Kim et al, 2019), we also investigated how autistic traits and sensory sensitivities are related to the relative weighting of prior and new information during a perceptual decision. Here, we empirically assess these theoretical models by employing Bayesian modelling of behavioural data captured in a task that manipulates uncertainty in priors and likelihoods (Körding & Wolpert, 2004; Vilares et al, 2012; Vilares & Kording, 2017). These studies have consistently demonstrated that people integrate prior and likelihood information in a qualitatively Bayesian-like fashion. While people often do not behave in a (quantitatively) perfectly Bayesian optimal way, the ways in which they deviate from this optimality can give us insights into the subjective information they have available (Tauber et al, 2017). Thus, because we are using Bayes as a framework to understand perceptual processes in autism, we do not test, nor do we compare the above-mentioned Bayesian models with non-Bayesian accounts of perceptual processes in autism. Our goal was to shed light on computational models of perception in autism.

## Methods

### Participants

We recruited a total of 80 adult participants (48 Neurotypicals and 32 participants who self-identified as having received a diagnosis of autism spectrum disorder). Recruitment was undertaken via Asperger’s Services Queensland, Autism Queensland, Mind and Hearts, The University of Queensland (UQ) online SONA system, and online advertisements. All participants (or their guardians) completed an online screening form. Participants were included in the study if they were between the ages of 18 – 35 years and had no history of neurological abnormalities. Participants were recruited initially if they reported having received a diagnosis of autism spectrum disorder from a clinician. Neurotypical participants had an additional inclusion criterion of no history of neurological abnormalities or psychological disorders and reported no current use of any medication acting on the nervous system. For group analysis, the autism spectrum (AS) group had 25 participants with a confirmed diagnosis of ASD, and the neurotypical (NT) group had age and gender matched 25 participants. For the dimensional analysis all 80 participants were included, this consisted of 48 neurotypical, 25 autistic and 7 ‘other’ participants who could not be confirmed as being on the autism spectrum using the ADOS. Thus, no participant’s data were excluded from the study. All participants provided written informed consent to participate in the study and were compensated at a rate of AUD20 per hour for their participation in the study. This study was approved by the Human Research Ethics Committee of The University of Queensland (Approval No.: 2019000119).

### Procedure

32 participants who reported having received a diagnosis of ASD from a clinician undertook a diagnostic interview with a clinical psychologist using the Autism Diagnostic Observation Schedule (ADOS) for Adults (Gotham, 2006; Hus & Lord, 2014), to confirm diagnosis and characterize symptom severity. The assessment lasted approximately 1-hour and was conducted on a separate day from the experimental sessions. Of the 32 participants, six individuals who scored below 3 on the ADOS were excluded from the autism spectrum group analysis. Additionally, one participant with a self-diagnosis of ASD was not available to complete the ADOS assessment. Thus, a total of 25 participants were confirmed to be on the autism spectrum and were included in the autism spectrum (AS) group (See Figure 1).

### Psychometric measures

All participants completed self-report questionnaires including the 50-item Autism Quotient (AQ) questionnaire (Baron-Cohen et al, 2001) and 93-item Sensory Processing Quotient (SPQ) Questionnaire (Tavassoli et al, 2014a) which were used to measure autistic traits and sensory sensitivities respectively. Participants also completed the Beck Anxiety Inventory (Beck et al, 1988) and Beck Depression Inventory (Beck et al, 1961).

### 2-Prior Coin Task (Vilares et al. 2012)

Participants engaged in a modified version of the visual decision-making task developed by Vilares and colleagues (2012), performed either in a 3T Magnetic Resonance Imaging scanner (AS = 28 and NT = 47) or outside of the scanner at a computer (AS = 4 and NT = 1). Participants were shown an image of a pond on a screen (Figure 2) and were told that someone was throwing a coin to the middle of the pond (i.e., the middle of the screen). Participants were told that they would see trials from two different coin-throwers and that one was better at throwing to the centre than the other. Unbeknownst to the participant, thrower A was more precise than thrower B (with order counterbalanced across participants), throwing the coin closer to the middle more often (narrow prior) than thrower B (wide prior). Before each block, participants were shown which thrower (Thrower ‘A’ or Thrower ‘B’) was throwing next. On each trial, participants were shown five blue dots representing the splashes that the coin made when falling into the pond. For each trial, participants were instructed to use a keyboard/button box to move a blue bar (“net”) horizontally across the screen to where they thought the coin had fallen on that trial. Next, participants moved a bar horizontally to rate how confident they were about their decision on a scale ranging from 0 (Guessing) – 100 (Confident). All these events were self-paced. The participant was then shown the true position of the coin, as a yellow dot, for 1500ms.

**Figure 2:**
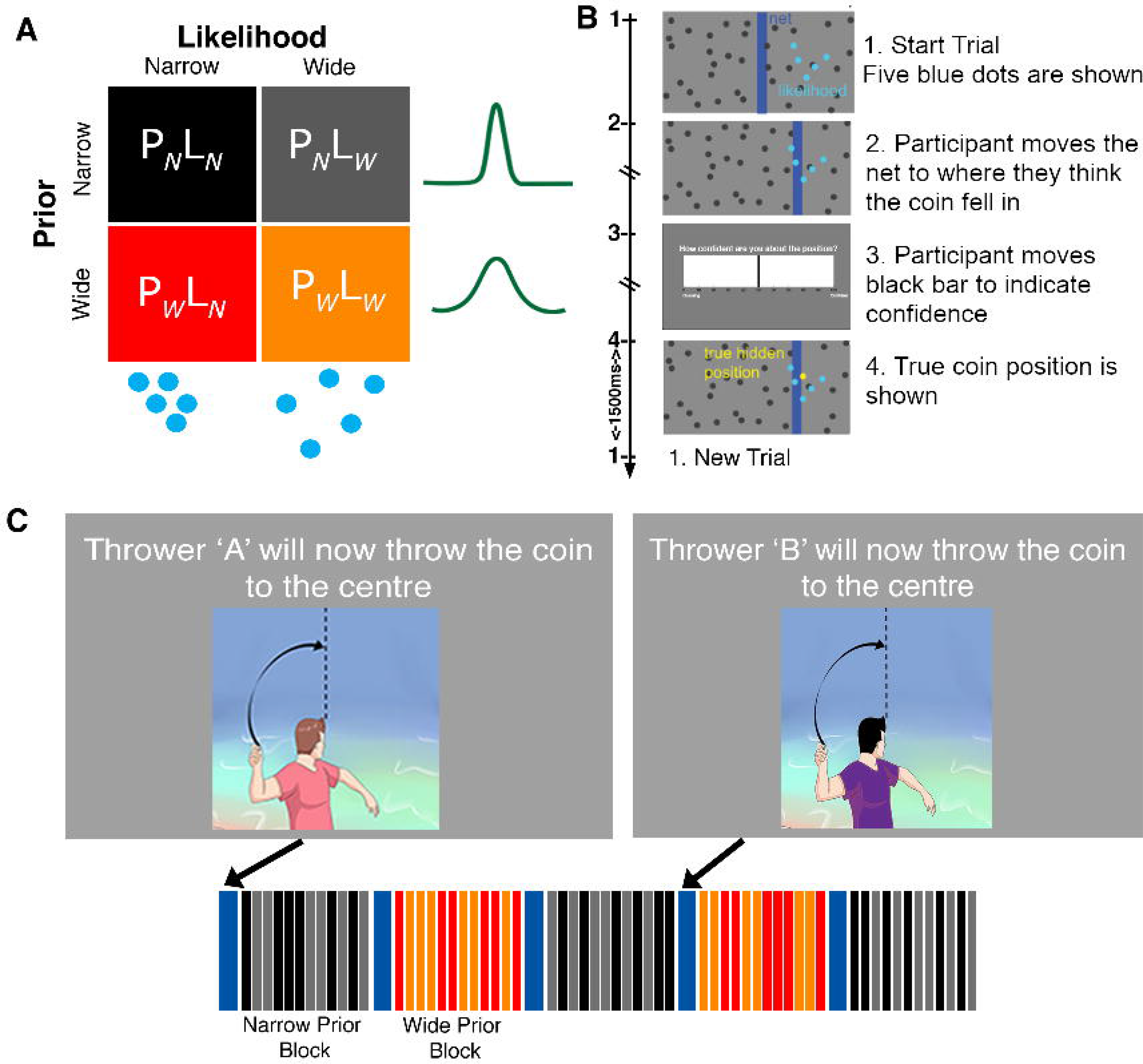
Coin Task Set up. A) Task design adapted from Vilares et al. (2012): the four conditions of the task - with two types of prior (P_N_ = narrow prior; P_W_= wide prior) and two types of likelihood (L_N_= narrow likelihood; L_W_= wide likelihood) uncertainty. B) The time course of a single trial: participants are asked to estimate the position where a coin fell given where the 5 splashes appeared. C) The trials were organized into 24 short blocks of 12 trials each with participants being told at the beginning of each block which thrower (“A” or “B”) will be throwing the coin (blue bar). Four types of trials are shown - Narrow Prior - Narrow Likelihood (P_N_L_N_, black); Narrow Prior - Wide Likelihood (P_N_L_W_, grey); Wide Prior - Narrow Likelihood (P_W_L_N_, red); and Wide Prior - Wide Likelihood (P_W_L_W_, orange)

The *prior variance* was manipulated across blocks as follows. The coin position was drawn from a Gaussian distribution in every trial, centred on the centre of the screen with a standard deviation that was either narrow (σ_PN_=2.5% of the screen width; Thrower A) or wide (σ_PW_=8.5% of the screen width; Thrower B) across blocks. The true prior variance was constant within a block, whereas the variance of the likelihood changed pseudo-randomly within a block. The variance of the splashes (i.e., the five blue dots) was the *true likelihood variance*. The spread of these dots could be narrow or wide, corresponding to narrow or wide likelihood variance, respectively. The position of the five dots on the x-axis of the screen was drawn from a Gaussian distribution with a mean corresponding to the true coin position and standard deviation either narrow (σ_LN_=6%) or wide (σ_LW_=15%). Where the variance of the five dots on the x-axis was narrow, the standard deviation of the dots on the y-axis was wide (σ=15%), and correspondingly narrow (σ =6%) on y-axis if wide on the x-axis. This was to ensure that the total area of the spread of the dots was uniform across both narrow and wide conditions and was the same area on the visual regions. However, since participants moved the bar only on the x-axis the true likelihood variance on each trial was calculated as the standard deviation on the x axis.

Thus, the experiment conformed to a 2x2 design with Prior (wide and narrow) by Likelihood (wide and narrow), consisting of 4 types of trials/conditions: *Narrow Prior - Narrow Likelihood* (P_N_L_N_, black); *Narrow Prior - Wide Likelihood* (P_N_L_w_, grey); *Wide Prior - Narrow Likelihood* (P_w_L_N_, red); and *Wide Prior - Wide Likelihood* (P_w_L_w_, orange).

**The practice task**: consisted of only 2 blocks (one per thrower/prior type, with 40 trials each thrower), taking between 10-15 minutes to complete.

**The main task:** consisted of 12 blocks per Prior, with each block having 12 trials from a single thrower (Thrower A or B) with 2 types of likelihoods (narrow/wide). Before the thrower changed participants were instructed as to which thrower, A or B, would be throwing next (5 seconds); See Figure 2C. In total, the main task consisted of 288 trials (72 trials per condition). The task duration was self-paced and took between 35 and 60 minutes to complete.

### Likelihood Only Task

After completing the 2-Prior Coin Task, participants completed a Likelihood Only Task outside of the scanner at a computer. Four participants in the AS group and one participant in the NT did not complete this task, as they did not wish to continue due to fatigue. The aim of this task was to estimate the participant’s perceived likelihood distribution, i.e., how the participants represent the centre of the dots on their own, without prior knowledge. Participants saw trials as in Figure 2B. However, participants did not report confidence on this task and the true coin position was always the centre of the splashes. Participants were simply instructed to move the net to where they thought the centre of the splashes was (i.e., the middle of the 5 blue dots). The true centre of the dots was shown in yellow as feedback to each participant on every trial. This task consisted of only one block with 144 trials from 2 types of likelihood (narrow/wide) and took between 15 - 20 minutes to complete. This task was conducted after the 2-Prior Task, as we expected that the participants would be biased towards the centre of the splashes if they performed the Likelihood Only Task first and thus would have a difficulty in learning the prior in the main 2-Prior Task that followed. In hindsight, however, we acknowledge that this caused the participants to carry over their priors (from the 2-Prior Task) to the Likelihood only tasks, hence defeating our purpose of having a no-prior task. Future studies may wish to consider administering the Likelihood-only task first or counterbalance the two tasks.

### Behavioural Analysis

From Bayes rule, we can obtain what would be the optimal estimate for the position of the coin on each trial of the coin-catching task (For detailed workings see (Körding & Wolpert, 2004; Vilares et al, 2012)):

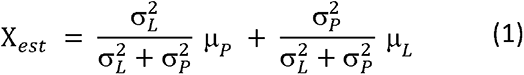

where X_est_ is the estimated position of the coin (i.e., participant responses on each trial). (σ^2^_P_, μ_P_) and (σ^2^_L_, μ_L_) represent the variance and mean of the prior (i.e., participant’s subjective model of where each thrower would throw the coin) and likelihood/sensory observation distributions (i.e., participant’s subjective measure of the five blue dots).

### Estimating likelihood vs. prior reliance

As described in Vilares et al. (2012), for each condition we can fit a linear regression to predict the participants’ estimated position of the coin for each trial (X_est_) as a function of the likelihood mean (μ_L_ here are the centre of the splashes). The slope of the regression line (the term 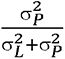) in equation (1) is the sensory weight (sw / likelihood reliance), which indicates how much the participant relies on the likelihood/sensory information (see Figure 3A). The closer the slope is to one, the more the participant relied on likelihood information (i.e., centre of five blue dots). If we assume that participants only rely on current and/or prior information (e.g., if we exclude random behaviour), then a slope closer to zero corresponds to relying more on the prior (i.e., centre of the screen), with anything in between indicating integration of the likelihood and the prior.

**Figure 3:**
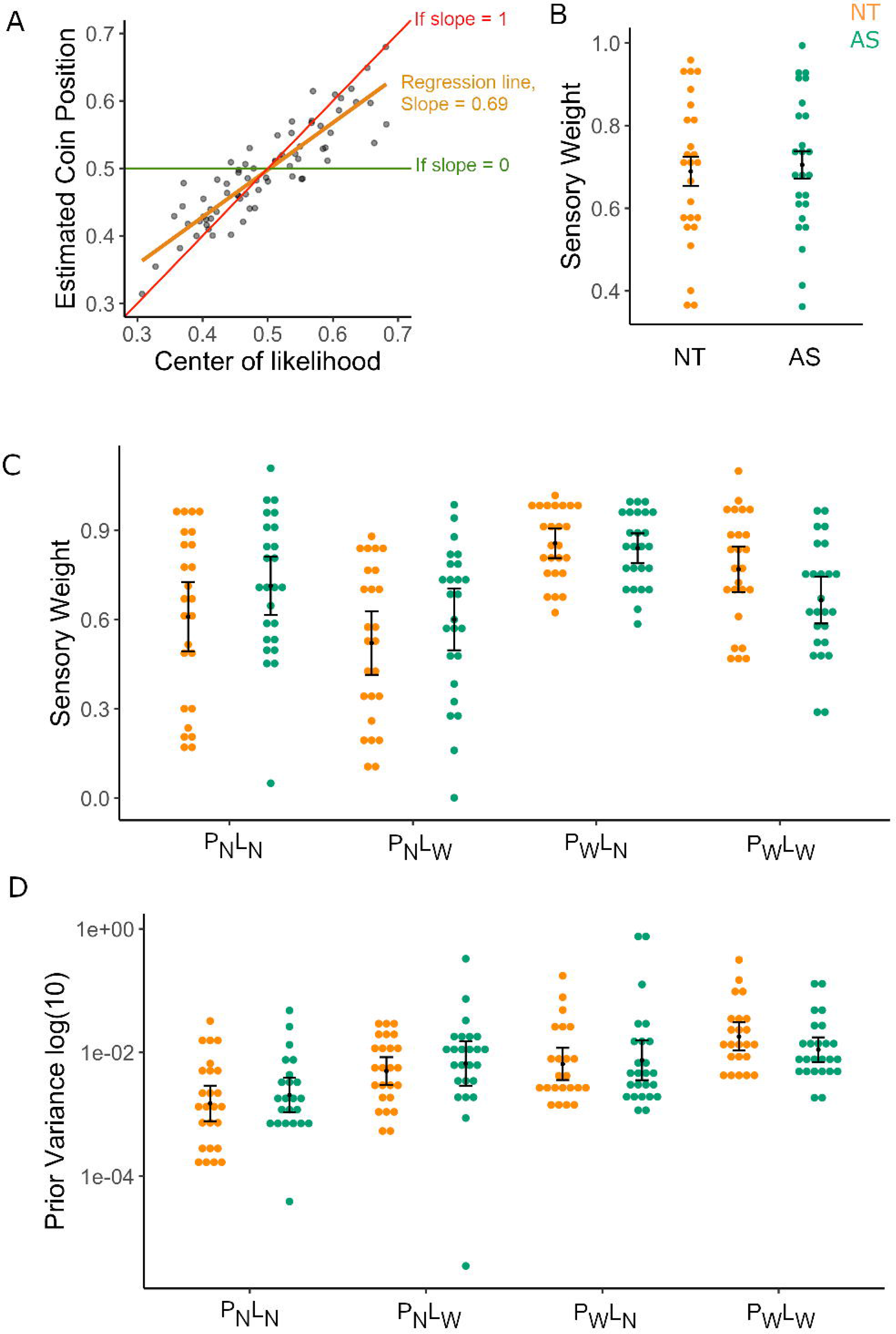
Likelihood vs. Prior Reliance in the 2-Prior Task. A) Sensory weight (likelihood reliance) for a participant is calculated by obtaining the slope of the regression (orange) between the true centre of the likelihood and participant’s estimates of the coin position for each condition. Slopes closer to 1 (red) indicate that participants are more reliant on likelihood information (mean of the splashes) while slopes closer to zero (green) indicate that the participant didn’t rely much on likelihood information, suggesting that they may have relied on prior information instead. Figure shows an example participant. Each dot corresponds to the response on a given trial. B) Sensory weights per group (Left, NT- orange; right, AS-green) averaged across the four conditions. No group differences were found in likelihood reliance in the Main (2-Prior) Coin Task. C) Average sensory weights per group, separated by condition. The AS group shows reduced context adjustment compared to NT. D) Estimated subjective prior variances, divided per group and condition. No evidence was found for group differences in subjective prior variance. Conditions: P_N_L_N_ = narrow prior and likelihood; P_N_L_W_ = narrow prior, wide likelihood; P_W_L_N_ = wide prior, narrow likelihood; P_W_L_W_ = wide prior and likelihood.

### Estimating a participant’s subjective prior mean

From equation (1), the intercept of the regression line (the term 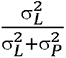) can be rearranged to calculate the prior mean acquired by the participant:

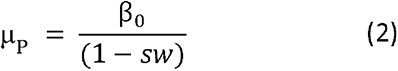

### Estimating participant’s trial by trial sensory weights

Equation (1) can be rearranged to obtain a trial-by-trial slope (i.e., sensory weight)

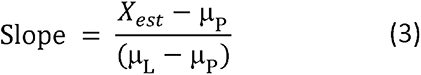

### Estimating participants’ subjective prior variance

Again, from equation (1), the slope or the sensory weight (sw) is equal to 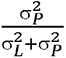. This can be rearranged to obtain the participant’s subjective prior:

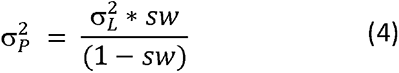

In equation 2, σ^2^_L_ can be assumed as the true or objective likelihood variance (given by the variance of the 5 blue dots). or can be estimated from the subjective likelihood variance (σ^2^_Ls_; described below in equation 3).

### Estimating a proxy for subjective likelihood variance

The variance of the participant’s estimates of the mean (μ_est_) relative to the true mean of the splashes (μ_L_) on the Likelihood Only task can be determined as a proxy for the participant’s subjective likelihood variance 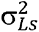 is as follows:

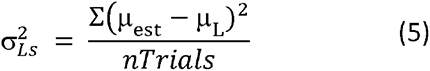

### Statistical Analysis

We aimed to understand if any of the current Bayesian models can explain sensory learning in autism. To test these models, we estimated the subjective prior and likelihood variance measures for each participant as described above.

For group analyses, we conducted 2x2x2 repeated measures analysis of variance for each measure of interest, with Prior (narrow vs wide) and Likelihood (narrow vs wide) and Group (AS vs NT) as factors. The different outcome variables analysed were: 1) Sensory weight or likelihood reliance; 2) estimation error (i.e., participants’ estimate minus the true position of the coin) for narrow and wide likelihood conditions as the outcome variables; 3) subjective prior variance for each condition, and 4) average confidence. Independent Samples T-tests were conducted for post-hoc pairwise comparisons. Where appropriate, Bayes factors (BF_01_) are also reported in support of evidence for the null hypothesis. For all group tests estimated marginal mean (EM Mean) and effect size (partial eta^2^/η_p_^2^) are reported.

For the dimensional analyses, we conducted bootstrapped Spearman rank correlations with AQ and SPQ scores with likelihood reliance and subjective prior variance for the main 2-prior task. Bootstrapped 95% confidence intervals for Spearman correlations reported are based on 1000 samples. For the Likelihood Only Task we excluded outliers based on Tukey’s 1.5 Interquartile Range. Based on this test 7 participants’ data (1NT and 3AS and 3 Other) were excluded.

In order to assess if continuum results were driven by group differences (AS vs NT vs other) we further, conducted multivariate analysis of covariance with AQ as covariate, Group as fixed factor and variable interest as the outcome variable to assess the interaction between group and AQ. Corrections for multiple comparisons are reported based on the Bonferroni correction procedure (*p*_bonf_) alongside the uncorrected *p* value. Bonferroni corrections are applied considering all statistical comparisons with a trait of interest (e.g., AQ score) within a task (i.e., within Likelihood Only or within 2-Prior Task). Statistical analysis was conducted in SPSS version 26 and R. Figures are presented using ggplot2 (Wickham, 2016) in RStudio.

## Results

### Participants

A total of 80 adults participated in the study. Demographic details and psychometric profiles are provided in Table 1. Group analysis was conducted on 25 participants confirmed to be on the autism spectrum (AS Group) and 25 age and gender matched neurotypical adults (NT group). The NT group showed lower autistic traits (*t* = -6.712, *p* = 2.161 x 10^-8^) and lower visual hypersensitivities (i.e., higher SPQ scores) than the AS group (*t* = 2.846, *p* = 0.007). The two groups showed no differences in anxiety (t = -1.466, p = 0.149) and depression (t = -1.980, p = 0.055). However, 12 participants in the AS group reported current anti-anxiety/ antidepressant use and 5 participants reported medication for ADHD. Neurotypicals reported no current medication use.

**Table 1 :** Demographic profiles and scores from self-administered psychometric scales

| Variable | Neurotypical (NT) Group<br>(*n* = 25) |  |  | Autism Spectrum (AS) group<br>(*n* = 25) |  |  | Total Sample<br>(*n* = 80)<br>(48 NT + 25 AS + 7 Other) |  |  |
| --- | --- | --- | --- | --- | --- | --- | --- | --- | --- |
|  | *M* | *SD* | Range | *M* | *SD* | Range | *M* | *SD* | Range |
| Age (years) | 23.96 | 4.82 | 18 - 35 | 25.48 | 6.501 | 18 - 35 | 24.69 | 5.16 | 18 - 35 |
| Sex at Birth (F/M/Intersex) | 15/10/0 |  |  | 14/10/1 |  |  | 37/42/1 |  |  |
| Gender (F/M/Other*) | 14/10/1 |  |  | 13/10/2 |  |  | 39/35/5 |  |  |
| Autism Quotient (AQ) | 19.44 | 7.25 | 4 - 30 | 35.44 | 7.42 | 21 - 46 | 26.15 | 10.24 | 4 - 46 |
| AQ Attention to detail | 4.64 | 2.66 | 1 - 10 | 6.40 | 2.45 | 1 - 10 | 5.52 | 2.68 | 1 - 10 |
| AQ Attention switching | 4.68 | 1.93 | 2 - 9 | 8.52 | 1.66 | 3 - 10 | 6.60 | 2.63 | 2 - 10 |
| AQ Social Skill | 3.88 | 2.29 | 0 - 7 | 7.32 | 2.17 | 3 - 10 | 5.60 | 2.81 | 0 - 10 |
| AQ Communication | 3.56 | 2.10 | 0 - 8 | 8.28 | 1.62 | 4 - 10 | 5.92 | 3.02 | 0 - 10 |
| AQ Imagination | 3.08 | 1.97 | 0 - 8 | 4.92 | 2.41 | 1 - 9 | 4.00 | 2.37 | 0 - 9 |
| Sensory Processing Quotient (SPQ) | 117.76 | 14.23 | 90 - 159 | 106.33 | 27.94 | 58 - 162 | 108.99 | 23.85 | 50 - 162 |
| SPQ Vision subscale | 28.96 | 3.82 | 23 - 38 | 24.44 | 6.96 | 13 - 44 | 25.94 | 5.99 | 13 - 44 |
| SPQ Hearing subscale | 28.32 | 4.99 | 19 - 38 | 26.92 | 5.07 | 19 - 37 | 27.62 | 5.03 | 19 - 38 |
| SPQ Smell subscale | 18.88 | 4.19 | 11 - 28 | 19.64 | 6.73 | 9 - 34 | 19.26 | 5.56 | 9 - 34 |
| SPQ Taste subscale | 20.12 | 4.10 | 14 - 29 | 16.68 | 8.29 | 1 - 33 | 18.40 | 6.70 | 1 - 33 |
| SPQ Touch subscale | 21.48 | 4.56 | 13 - 29 | 17.36 | 6.70 | 3 - 32 | 19.42 | 6.03 | 3 - 32 |
| Beck Anxiety Score | 13.68 | 11.60 | 0 - 55 | 18.72 | 12.68 | 0 - 44 | 14.50 | 11.67 | 0 - 55 |
| Beck Depression Score |  |  |  | 16.04 | 13.61 | 0 - 44 | 11.09 | 9.89 | 0 - 44 |
| Antidepressant use(Y/N) | 0/25 |  |  | 12/13 |  |  | 17/63 |  |  |
| ADHD medication use (Y/N) | 0/25 |  |  | 6/19 |  |  | 8/72 |  |  |
| Autism Diagnostic Observation Schedule (ADOS) | N/A |  |  | 6.64 | 1.78 | 4 - 10 | 5.87 | 2.26 | 2 - 10 |

Note: \*Other Genders include – female to male transgender (1NT & 2 AS)

Dimensional analysis pooled all 80 participants together (47 NT + 32 self-reported ASD) aligning them on Autism Quotient (AQ) scores. AQ scores were negatively correlated with SPQ vision scores [*r* = -0.463, p = 1.6 x 10^-5^] indicating greater hypersensitivities for higher AQ scores as well as increased anxiety [*r* = 0.422, p = 9.8 x 10^-5^] and depression [*r* = 0.349, p = 0.002] with AQ.

We first report our findings on group differences and then turn to the analysis of autistic traits by aligning all participants on a continuum using autism quotient scores.

### No group differences in task performance

In the practice task, a 2x2x2 ANOVA of prior means (see methods equation 2 for calculation), showed no group differences in their subjective prior mean [NT group mean = 0.560; AS group mean = 0.526; *F* = 0.630, *p* = 0.431]. A 2x2x2 ANOVA of estimation errors demonstrated no differences in overall accuracy [*F* = 1.450, *p* = 0.234]. This indicated that all participants regardless of group, had acquired the centre of the screen as the prior mean (as instructed). Furthermore, there were no group differences in overall task performance.

Further, for the main 2-prior task, to establish task performance, we conducted a 2x2x2 ANOVA for estimation errors. We observed a main effect of Prior, driven by P_W_>P_N_ [M = 8.7 x 10^-3^, p = 1.067 x 10^-8^]; a main effect of Likelihood, driven by L_W_>L_N_ [ M = 0.025, *p* = 3.55 x 10^-24^] and a Prior*Likelihood interaction [*F* = 19.276, *p* = 6.20 x 10^-5^]. This indicated that the prior and likelihood uncertainty manipulations functioned as expected, with higher uncertainty in the prior or likelihood leading to more estimation errors. We found no Group interactions with Prior or Likelihood factors for estimation errors, indicating that the AS and NT group showed no differences in performance (i.e., estimation errors, See sup. Table S2 and sup Figure 4A).

**Figure 4:**
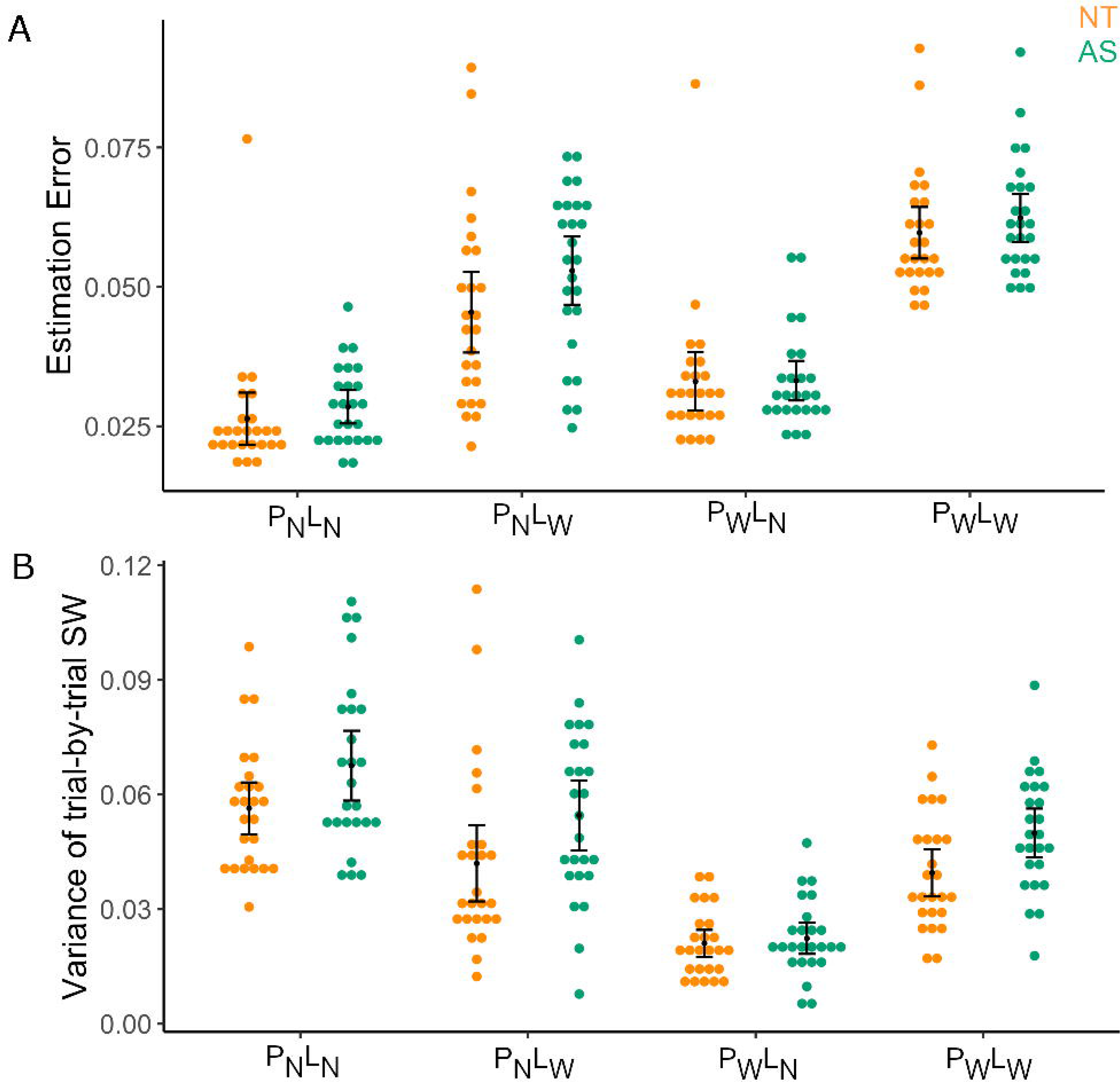
Performance in the 2-Prior Task. A) Estimation Errors in the 2-Prior Task shows no difference in performance between groups by condition B) Variance of trial-by-trial sensory weights reveal a main effect of Group driven by higher variability in the AS group. (NT: Neurotypical - orange; AS: autism spectrum group -green).

We confirmed that priors were learnt in the main task and that participants were able to discriminate conditions as demonstrated by the predicted sensory weights (see Figure 3C) showing an effect of Prior [η_p_^2^ = 0.416, *F* = 34.129, *p* = 4.35 x 10-7] and Likelihood [η_p_^2^ = 0.521, *F* = 53.304, *p* = 3.24 x 10-9] regardless of group whereby the sensory weight is higher for more reliable likelihood information and lower for more reliable priors.

Groups also showed no differences in their confidence reports for individual conditions (See sup. Table S1.4), but we did observe a Group*Likelihood interaction [η_p_^2^ = 0.160, p = 0.004]. This interaction is driven by the AS group showing a smaller difference [*M* = -2.176, *SD* = 6.812] than the NT group [*M* = -9.618, *SD* = 10.223] in their confidence reports in wide vs narrow likelihood (i.e., L_W_ – L_N_) conditions (See Sup Figure S1).

### No significant difference in likelihood reliance between groups

We first aimed to establish whether the AS group gave more weight to new (likelihood) information than prior information than the NT group (See Methods; Figure 3A) using data from the main 2-Prior Task. Contrary to the hypothesis of increased likelihood reliance in AS, as posited by the theoretical work of Brock (2012) and Pellicano & Burr (2012), we found evidence for no difference between the groups in overall likelihood reliance, (i.e., sensory weights averaged across conditions) [*t* = -0.325, BF**_01_** = 4.524, *p* = 0.747; Figure 3B].

### AS group shows less context adjustment in sensory weights

Looking within conditions in the 2-Prior Task, a repeated-measures ANOVA (see methods) of sensory weights revealed a significant Group*Prior [Effect Size η_p_^2^ = 0.123, *p* = 0.013], but no Group*Likelihood [η_p_^2^ = 0.058, *p* = 0.091] interaction or Group*Prior*Likelihood [η_p_^2^ = 0.026, p = 0.260] interaction (see Figure 3C). Post-hoc tests revealed the Group*Prior interaction effect was driven by the NT group showing a larger difference in sensory weights for wide prior vs. narrow prior (i.e., P_w_ - P_N_), [*M* = 0.495, *SD* = 0.450], compared to the AS group [*M* = 0.191, *SD* = 0.377], who presented less shift in sensory weights across contexts.

Between-group pairwise comparisons revealed no statistically significant differences between groups for individual conditions (see Supp. Table S1 for statistics for each condition).

### Intact priors in autism: no evidence for differences in subjective priors between groups

Participants’ behavioural responses on the 2-Prior Task were modelled using a Bayesian approach to determine the subjective prior variance (see Methods for details). When investigating prior variance, we did not observe any significant Main effect of Group [η_p_^2^ = 0.029, *p* = 0.250], Group*Prior [η_p_^2^ = 0.008, *p* = 0.545], Group*Likelihood [η_p_^2^ = 0.039, *p* = 0.176] or a Group*Prior*Likelihood interaction [η_p_^2^ = 0.078, *p* = 0.054] (see Figure 3D). We find no evidence for differences between groups in individual conditions (see Supplemental Section S1.3).

### AS group shows more variability in trial-by-trial sensory weights

We further investigated variability in sensory weight by obtaining the variance in trial-by-trial sensory weights (See Equation 3). We found a main effect of Group [η_p_^2^ = 0.084, *p* = 0.042] but no further group interactions with prior or likelihood (See Supp. Table S8). This main effect of group was driven by higher variability in the AS group [*M* = 0.049, 95% CI = 0.043 to 0.055] compared to the NT group [*M* = 0.040, 95% CI = 0.034 to 0.046]. Further, individual conditions revealed differences to be in P_N_L_N_, P_N_L_W_, P_W_L_W_ conditions (See Figure 4B and Suppl. Table S9 for statistics).

### The AS group showed a stronger bias to the centre of the screen in the Likelihood Only Task

A subsample of 21 AS and 24 matched-NT also completed a likelihood only manipulation task after the main 2-Prior Coin Task, in which the prior manipulation was removed, and participants’ task was to find the centre of the cloud of dots (See methods). This Likelihood Only task was designed to obtain an estimate of the variance of participants’ sensory observations (i.e., the likelihood variance), in the absence of a given prior.

However, results from this task revealed that the AS group’s estimates of the centre of the splashes was shifted toward the middle of the screen, as shown by significantly lower sensory weights than those found in the matched-NT group, for both the Narrow [Mann-Whitney U = 130.00, *p* = 0.006] and the Wide [Mann-Whitney U = 128.00, *p* = 0.005] likelihood conditions (See Figure 5B). However, this did not manifest as a significant difference in accuracy (i.e., mean estimation error) between the groups in this task (see Figure 5A). This result may reflect a carryover bias in the Likelihood Only task from the 2-prior task. We therefore did not use estimation error variance as a measure of subjective likelihood variance for group differences.

**Figure 5:**
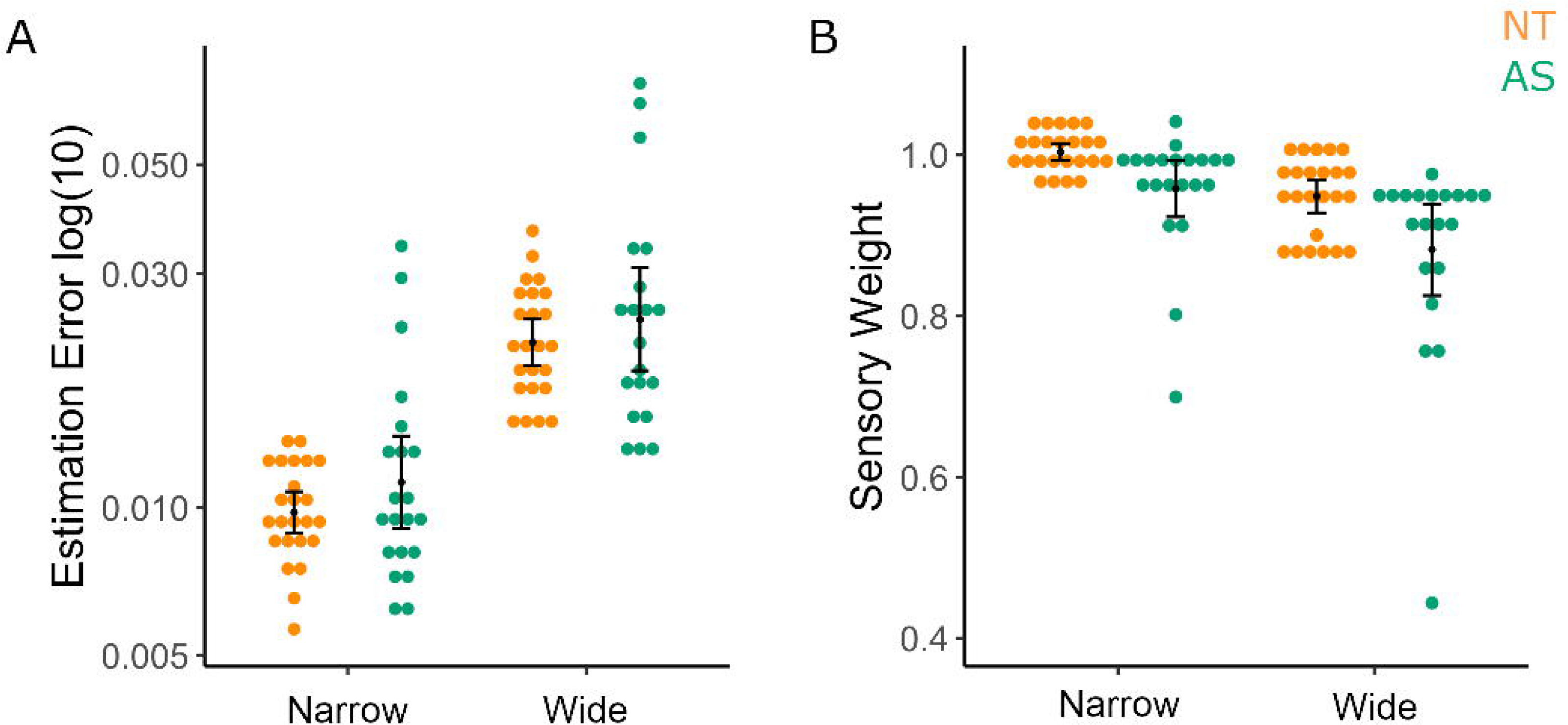
Likelihood Only Task results. reveal that the AS group has high variability (i.e., were further from estimating the true centre of the splashes) but show no difference in A) estimation errors compared to the NT group for either Narrow or Wide likelihood variance conditions. B) AS group shows lower sensory weights compared to NT for Narrow and Wide likelihood conditions which may indicate a carryover bias from the main 2-Prior task. (Left, NT- orange; right, AS-green).

### No effects of medication on group findings

Seventeen participants in the AS group reported taking antidepressant medication and 8 participants reported medication use for attention deficit hyperactivity disorder (ADHD). In contrast, none of the NT participants reported any medication use, which may be a confounding factor in our group analyses. We undertook a control analysis to identify whether medication use was a significant predictor of the variables of interest within the AS group. To that end, we conducted a regression analysis with variables of interest as the outcome variable. We found no relationship between antidepressant or ADHD medication use difference in sensory weights (P_W_ – P_N_) (*F* = 0.081, *p* = 0.922) in the main 2-Prior Task. Further, our estimates of sensory weights for the Likelihood Only task showed no relationships with medication use for narrow (*F* = 0.625, *p* = 0.543) or wide (*F* = 0.431, *p* = 0.655) likelihood conditions. Of course, with the relatively small numbers of participants reporting medication use here, we cannot exclude the possibility that there was a subtle but undetected effect of medication on the behavioural findings.

### Autistic traits are not correlated with accuracy or prior variance but are negatively correlated with confidence in the 2-Prior task

We undertook a dimensional approach by aligning the participants on a continuum using the dimensions of both their Autism Quotient (AQ) scores, which quantify autistic traits, and their Sensory Processing Quotient (SPQ) scores, which assess sensory sensitivities. In the main 2-Prior task, we found no associations that survived multiple comparisons, between AQ and SPQ scores and likelihood reliance, subjective prior variance, or accuracy (estimation error) in the pooled sample (AS and NT combined, see supplements Table S10). However, despite behavioural accuracy measures showing no significant relationships with AQ scores, we observed a lower mean confidence with increased AQ scores [*r* = -0.339, *p* = 0.002, *p_bonf_* = 0.034]. An ANCOVA revealed within group (AS vs NT) differences in their correlations with confidence and AQ, indicating pooled samples should be interpreted with caution. However, further within-group analysis showed that neither AS group [r = -.103 p = 0.575] or NT [r = -.189, p = 0.197] were driving the correlation in the pooled sample. We also conducted a within-AS group correlational analysis with the ADOS scores (which describe symptom severity within the autism group) but found no relationship with likelihood reliance or subjective prior variance (See supplements Table S17).

### Autistic traits but not sensory sensitivities are negatively associated with subjective likelihood variance in the Likelihood Only Task

We did not find a significant relationship between sensory weights in the Likelihood Only task and AQ for the narrow [*r* = -0.154, *p* = 0.188] or wide [*r* = -0.137, *p* = 0.242] likelihood conditions. We nonetheless excluded significant outliers by sensory weights in the Likelihood Only task (see methods). After excluding outliers, there was no significant relationship between sensory weights and AQ for the narrow [*r* = -0.147, *p* = 0.236] or wide [*r* = -0.140, *p* = 0.259] likelihood conditions.

Estimation error (performance) showed a negative correlation with AQ only in the wide likelihood condition [*r* = -0.377, *p =* 0.003*, p_bonf_* = 0.012; Figure 6B]. Further, within-group analysis revealed the NT participants to show a negative correlation of AQ with estimation error [*r* = -0.363, *p* = 0.007, *p_bonf_* = 0.028] in the wide likelihood condition, but not within the AS group [*r* = -0.158, *p* = 0.560].

**Figure 6:**
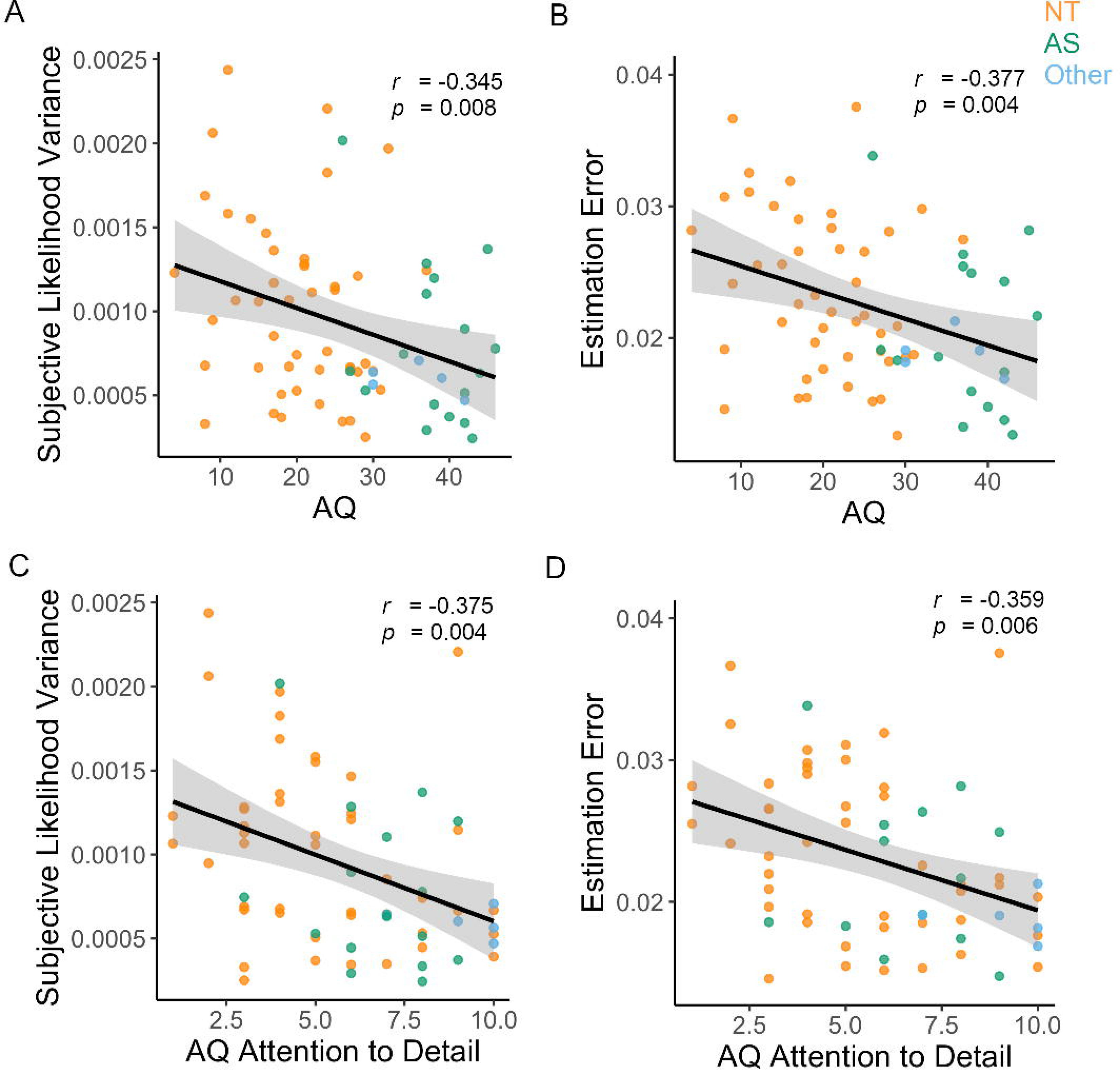
Autistic trait behaviour in the Likelihood Only Task. The Likelihood Only Task reveals negative correlations in the Wide Likelihood Condition between autism quotient (AQ) scores and A) subjective likelihood variance and B) Estimation error. An AQ subscale analysis revealed Attention to Detail to be negatively correlated with C) subjective likelihood variance and D) Estimation error.

We found a significant negative association [*r* = -0.326, *p =* 0.007*, p_bonf_* = 0.028; Figure 6A] between AQ scores and wide subjective likelihood variance (i.e., estimation error variance; See methods), but not with narrow subjective likelihood variance [*r* = 0.062, *p* = 0.618] in the pooled sample. This suggests that as autistic traits increase the precision of sensory observations increase which is apparent only in the wide likelihood condition.

Further, to check if groups could be pooled, using an ANCOVA we tested whether the AQ showed group (NT vs AS) differences in slopes (correlation coefficient) with wide likelihood variance on the Likelihood Only task [*F* = 3.107, *p* = 0.033; table S12]. This indicated that NT and AS groups showed differences in their correlations with AQ and could not be pooled. Within-group associations with AQ and subjective likelihood variance did not approach significance when corrected for multiple comparisons (See Supplements Table S11).

Further an AQ subscale analysis found that “*Attention to Detail*” negatively correlated with subjective likelihood variance [*r* = -0.375, *p* = 0.002, *p_bonf_* = 0.008; Figure 6C] and estimation error [*r* = -0.359, *p =* 0.003, *p_bonf_* = 0.012; Figure 6D] in the Wide Likelihood condition only in the Likelihood Only Task (Supplements Table S14). Once again, an ANCOVA revealed that within group correlations are significantly different between AS and NT for both estimation error [*F* = 3.965, *p* = 0.012] and likelihood variance [*F* = 3.337, *p* = 0.025] in the Wide Likelihood condition, suggesting caution in drawing conclusions on the pooled sample. Within-group analysis revealed the NT participants to show a negative correlation of Attention to Detail with subjective likelihood variance [*r* = -0.329, *p* = 0.026] and estimation error [*r* = -0.312, *p* = 0.035] in the wide likelihood condition. We did not observe significant correlations within the AS group for subjective likelihood variance [*r* = -0.194, *p* = 0.471] or estimation error [*r* = -0.184, *p* = 0.496].

## Discussion

In this study, we aimed to test whether individuals with AS showed greater likelihood reliance than NT individuals, consistent with current models of autistic perception, as well as to investigate whether this reliance aligned on a continuum of autistic traits and sensory sensitivities. To our knowledge, previous studies have not directly compared the relative weighting of sensory (likelihood) and contextual (prior) information in AS. The hypo-prior (Pellicano & Burr, 2012) and precise likelihood models (Brock, 2012) postulate an increased likelihood reliance. While our AS group did show significantly higher hypersensitivity (i.e., lower SPQ scores) than the NT group, contrary to both models, we found no difference in overall weighting of likelihood information relative to prior in the AS group (vs. NT). However, greater variability in trial-by-trial sensory weights in the autistic group suggests underlying difficulties in learning context. These findings are directly in line with a study that showed increased behavioural variability, but intact priors and integration in a visual task in ASD (Noel et al, 2020). In addition, we found that subjective likelihood variance decreased as autistic trait increased, which lends support to Brock (2012) precise likelihood model.

While task performance did not show any differences between groups, modelling of likelihood reliance (i.e., sensory weights) revealed a poorer adjustment across contexts in the AS group. Specifically, the AS group showed lower sensory weights compared to the NT group under a high uncertain (Wide Prior) context. This suggests a deficit in precision updating, and a higher reliance on prior (relative to likelihood) in the AS group when uncertainty is high. While Brock and Pellicano models are unable to explain a higher reliance on prior information, an alternate model, the HIPPEA model (Van de Cruys et al, 2014) predicts higher prior precision (which in turn may lead to greater reliance on prior). The high, inflexible precision of prediction errors (HIPPEA) model, on the other hand, argues that prediction error weighting is less flexibly adjusted in individuals with AS, particularly across different contexts (Van de Cruys et al, 2014). The HIPPEA model can explain instances in which both likelihood and prior variance can be of increased precision which may be a result of prediction error weighting. This deficit in prediction error weighting can explain some key diagnostic symptoms of AS, such as altered perceptual processing and resistance to change, as well as social differences relative to neurotypicals.

The HIPPEA model also makes specific predictions about alterations in prediction errors. Further research employing a paradigm specifically designed to investigate prediction errors would be needed to understand if the HIPPEA model may explain this reduction in context updating in AS. Reductions in contextual adjustment using social priors, such as imitating motor movements in children (Amoruso et al, 2019) and adults (Chambon et al, 2017) on the autism spectrum, suggest that our findings of alterations in contextual adjustment may extend to more complex perceptual processes and underpin core diagnostic symptoms of an ASD. However, Mottron (2019);(2014), cautions against generalizing descriptive findings, particularly as autistic perception can be both highly domain- (i.e. visual/auditory) and individual-specific.

In the ‘Likelihood Only’ task, the AS group showed greater reliance on the prior learnt in the 2-prior task (indicated by lower sensory weights than the NT group). This suggests over-reliance on a pre-learned prior in the AS group, or prior rigidity, when switching contexts (and tasks) that suggest “insistence on sameness” behaviours typically seen in AS. Hypersensitivities have been shown to have a mediating effect on insistence on sameness in autistic children (Black et al, 2017; Lidstone et al, 2014; Wigham et al, 2015), indicating that increased hypersensitivities relate to increased anxiety, which may in turn manifest in an insistence on sameness. Thus, in the future, it would be beneficial to measure other dimensions of restricted and repetitive behaviours, and also to investigate sensory learning in autism with low vs high anxiety groups to understand the relative contributions of these factors in perceptual decision-making.

While we did not find group differences in accuracy or confidence measures in the 2-Prior Task, we did find a negative correlation between autism traits and confidence reports. Retrospective confidence refers to an ability to judge the accuracy of one’s decisions (Mamassian, 2016). Bayesian theories of confidence judgements posit that they reflect the subjective probability that a decision is correct given the evidence (Pouget et al, 2016). However, we did not observe this expected relationship of higher accuracy and confidence in our task. Other non-Bayesian accounts of confidence judgements suggest that confidence reports reflect the subjective probability that an observer made the best possible decision (Adler & Ma, 2018; Li & Ma, 2020). Furthermore, Song et al (2011), finds that confidence reporting was separable from objective performance across tasks. It is unclear from our task if we are observing specific disruptions in metacognition as autism traits increase, or simply a general tendency to be less confident in sensorimotor decisions; for which we would need a secondary task to compare confidence reports and behaviour.

We further aimed to empirically disentangle the current Bayesian theories for perceptual alterations in autism that predict either increased precision in sensory information (likelihood) or wider priors. When aligned on a continuum, we observed negative correlations with AQ and likelihood variance as well as estimation error (but not with sensory weights), indicating that not only did sensory precision increase with autistic traits but so did accuracy albeit in the wide likelihood condition only. This observation of an association between likelihood variance and AQ in wide likelihood condition indicates the need for investigating autistic perception under varying degrees of uncertainty in future studies. Nonetheless, together with a lack of a significant correlation between AQ and subjective prior variance, is in line with previous findings in neurotypicals with autistic traits (Karvelis et al, 2018), providing support for the Precise Likelihood Model (Brock, 2012) when aligned on a continuum. We further found that this relationship was driven by the AQ subscale – ‘Attention to Detail’, which is a common perceptual enhancement in autism, often identified as a cognitive style (Baron-Cohen et al, 2009). However, the presence of a diagnosis confounds these findings, and we found the relationship between precision and AQ, attention to detail to be driven by neurotypical participants. A larger sample of autistic adults would be necessary to understand if increased precision can explain increased attention to detail in autistic adults.

Our study has several limitations. In this study we assumed that participants had acquired a prior by the end of the Main 2-Prior Task. While we demonstrate that participants showed effects of prior and likelihood in their sensory weights it may be possible that some participants had not yet fully learnt the experimental variance. Additionally, our version of the Coin Task had short blocks with multiple switches between throwers (priors), compared to Vilares (2012) et al., which had longer blocks but less block repetitions (just two block repetitions per thrower). This increases the difficulty of the task and may reduce sensitivity to detect differences in behaviour and prior variance between groups. In the Likelihood Only Task, we observed a bias towards the mean of the prior in the AS group, which did not allow us to make inferences on the likelihood variance. Future studies would benefit from starting with the Likelihood Only Task or counterbalancing the tasks across participants. We also did not model motor response noise in this task (i.e., where the participants put the bar may be different to where they intended it to be), which may account for observed group differences particularly within conditions.

Further, the autism spectrum is heterogeneous, and our AS sample was limited to participants who were able to read and provide consent on their own. This inevitably excluded participants on the autism spectrum who are non-verbal or have intellectual disabilities. This limits the generalizability of our findings as evidence for a global theory of autistic perception.  An additional consideration is that we have not investigated cognitive abilities such as verbal reasoning in participants, which may result in group differences due to differences in understanding task requirements. Future studies may benefit from accounting for cognitive abilities. It is also important to note that more than 50% of our AS group reported taking antidepressant medication and 25% reported ADHD medication use, which is a significant confounding factor in group findings. Moreover, while we hypothesized that visual sensory sensitivities as measured by SPQ scores would be associated with behavioural measures on the coin task over autistic trait scores, we did not find evidence for this. Thus, it remains unclear if narrow priors and narrow likelihood can explain hypersensitivities. In addition, our findings provide empirical evidence for increased precision in likelihood variance, although a larger sample would be needed to confirm these findings and understand the relative contributions of dimensional aspects of autism. It may well be that the heterogeneous nature of the autism spectrum itself explains the seemingly contradictory findings that take turns in supporting these alternative models in literature.

In conclusion, our findings provide evidence for intact priors in sensory perception in individuals on the autism spectrum and for the contextual nature of autistic perception that may explain behavioural differences in uncertain worlds.

## Data Availability

The de-identified data for this study has been made available at the following link on UQ e-space: https://doi.org/10.14264/uql.2020.981

## Acknowledgements

Radhika Tanksale for conducting ADOS assessments and Prof Kate Sofronoff for valuable advice on the project. All participants for their valuable time. The University of Queensland Research Training Programme for funding to RR. The University of Queensland Fellowship (2016000071) to MIG. Australian Research Council Centre of Excellence for Integrative Brain Function (ARC Centre Grant CE140100007) to JBM and MIG.

## Supplementary Material

### 1.0 Group Results

We conducted 2x2x2 repeated measures analysis of variance for each measure of interest, with prior (Narrow vs Wide) and Likelihood (Narrow vs Wide) and Group (AS vs NT) as Factors. We also report independent samples t-test comparisons for individual conditions as well.

F- Statistic (F), Effect size (partial eta^2^ / η_p_^2^), Bayes Factor with evidence for the null (BF_01_), t-statistic (t) and p-values (p) are shown.

### 1.1 2-Prior Task: Likelihood Reliance (i.e., sensory weight)

**Table S2:** Independent samples comparison of sensory weights by condition

| Outcome Variables | Group | Mean (M) | Bayes Factor ($BF_{01}$) | t-statistic | p-value |
| --- | --- | --- | --- | --- | --- |
| $P_N L_N$ | NT | 0.610 | 1.975 | 1.407 | 0.166 |
|  | AS | 0.714 |  |  |  |
| $P_N L_W$ | NT | 0.521 | 2.747 | 1.108 | 0.274 |
|  | AS | 0.601 |  |  |  |
| $P_W L_N$ | NT | 0.857 | 4.241 | -0.499 | 0.620 |
|  | AS | 0.840 |  |  |  |
| $P_W L_W$ | NT | 0.769 | 0.921 | -1.939 | 0.058 |
|  | AS | 0.666 |  |  |  |

### 1.2 2-Prior Task: Performance (Estimation Error)

**Table S3:** Results from repeated-measures ANOVA for estimation error

| Estimation Error | F-statistic | Effect Size ($\eta_p^2$) | p-value |
| --- | --- | --- | --- |
| Main Effect Group | 1.206 | 0.025 | 0.278 |
| Group*Prior | 1.772 | 0.036 | 0.189 |
| Group*Likelihood | 2.306 | 0.046 | 0.135 |
| Group*Prior*Likelihood | 0.971 | 0.020 | 0.329 |

**Table S4:** Between group effects for estimation error

| Outcome Variables | Group | M | $BF_{01}$ | t | p |
| --- | --- | --- | --- | --- | --- |
| $P_N L_N$ | NT | 0.026 | 3.574 | 0.795 | 0.430 |
|  | AS | 0.028 |  |  |  |
| $P_N L_W$ | NT | 0.045 | 1.486 | 1.624 | 0.111 |
|  | AS | 0.053 |  |  |  |
| $P_W L_N$ | NT | 0.033 | 4.739 | 0.045 | 0.965 |
|  | AS | 0.033 |  |  |  |
| $P_W L_W$ | NT | 0.060 | 3.372 | 0.874 | 0.387 |
|  | AS | 0.062 |  |  |  |

### 1.3 2-Prior Task: Subjective Prior Variance (calculated with true likelihood as set by the task)

**Table S5:** Results from repeated-measures ANOVA for prior variance

| Estimation Error | *F* | $\eta_p^2$ | *p* |
| --- | --- | --- | --- |
| Main Effect Group | 1.355 | 0.029 | 0.250 |
| Group*Prior | 0.372 | 0.008 | 0.545 |
| Group*Likelihood | 1.888 | 0 | 0.887 |
| Group*Prior*Likelihood | 3.914 | 0.078 | 0.054 |

**Table S6:** Between group effects on the subjective prior variance

| Outcome Variables | Group | *M* | *BF*<sub>01</sub> | *t* | *p* |
| --- | --- | --- | --- | --- | --- |
| P<sub>N</sub>L<sub>N</sub> | NT | 0.004 | 4.455 | -0.374 | 0.969 |
|  | AS | 0.003 |  |  |  |
| P<sub>N</sub>L<sub>W</sub> | NT | 0.009 | 2.544 | 1.184 | 0.250 |
|  | AS | 0.025 |  |  |  |
| P<sub>W</sub>L<sub>N</sub> | NT | 0.010 | 2.292 | 1.273 | 0.163 |
|  | AS | 0.072 |  |  |  |
| P<sub>W</sub>L<sub>W</sub> | NT | 0.039 | 3.158 | -0.945 | 0.314 |
|  | AS | 0.022 |  |  |  |

### 1.4 2-Prior Task: Confidence

**Table S7:** Results from repeated-measures ANOVA for confidence

| Estimation Error | *F* | $\eta_p^2$ | *p* |
| --- | --- | --- | --- |
| Main Effect Group | 2.381 | 0.047 | 0.129 |
| Group*Prior | 2.200 | 0.044 | 0.145 |
| Group*Likelihood | 9.172 | 0.160 | 0.004* |
| Group*Prior*Likelihood | 0.552 | 0.011 | 0.461 |

**Table S8:**
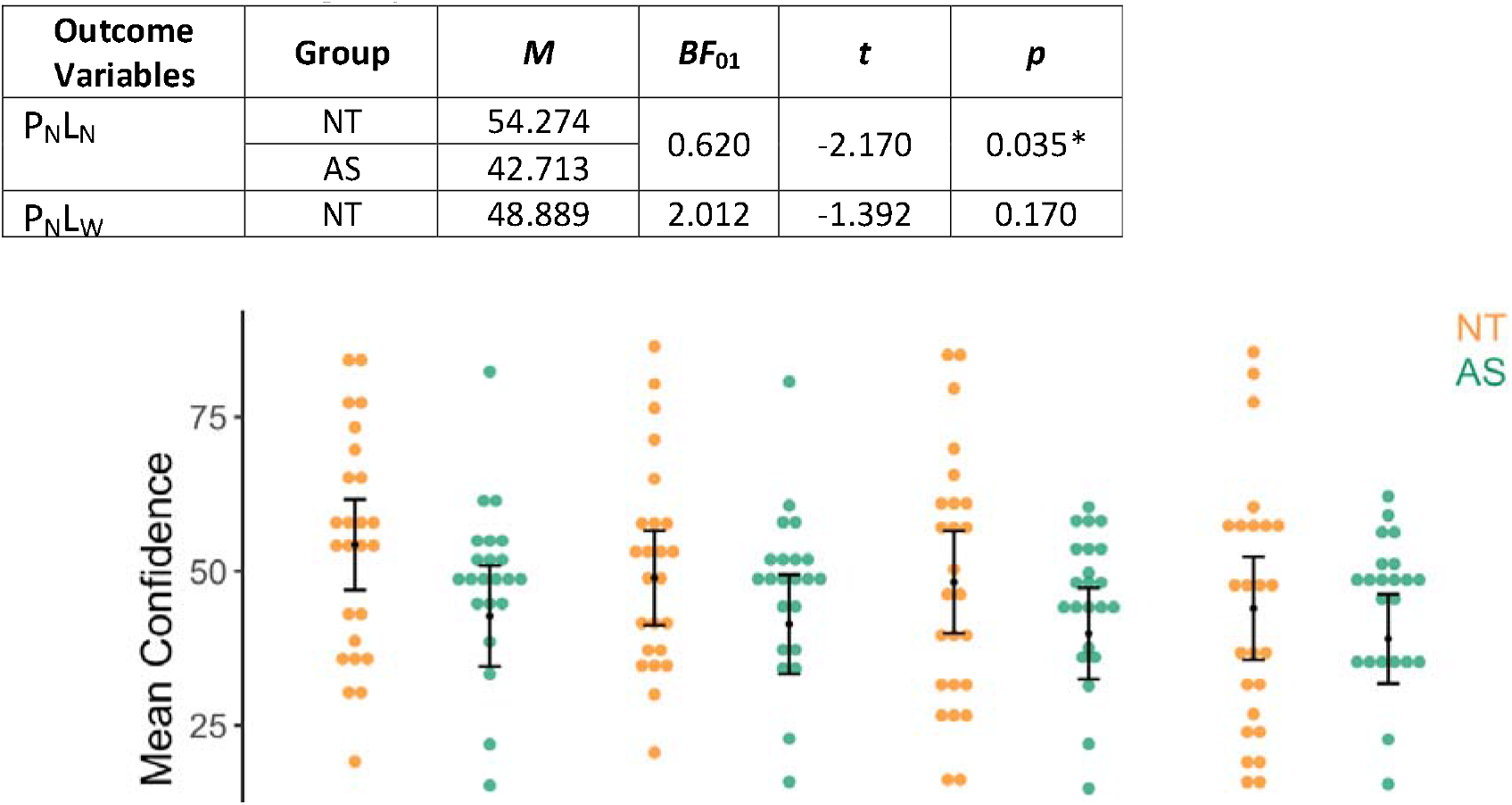
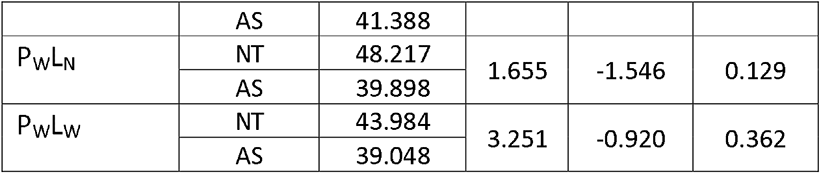
Between group effects on confidence

| Outcome Variables | Group | *M* | *BF*<sub>01</sub> | *t* | *p* |
| --- | --- | --- | --- | --- | --- |
| P<sub>N</sub>L<sub>N</sub> | NT | 54.274 | 0.620 | -2.170 | 0.035* |
|  | AS | 42.713 |  |  |  |
| P<sub>N</sub>L<sub>W</sub> | NT | 48.889 | 2.012 | -1.392 | 0.170 |
|  | AS | 41.388 |  |  |  |
| P<sub>W</sub>L<sub>N</sub> | NT | 48.217 | 1.655 | -1.546 | 0.129 |
|  | AS | 39.898 |  |  |  |
| P<sub>W</sub>L<sub>W</sub> | NT | 43.984 | 3.251 | -0.920 | 0.362 |
|  | AS | 39.048 |  |  |  |

### 1.5 2-Prior Task: Trial-by-trial sensory weights

**Table S9:** Results from repeated-measures ANOVA for trial-by-trial sensory weights

| Estimation Error | *F* | $\eta_p^2$ | *p* |
| --- | --- | --- | --- |
| Main Effect Group | 4.381 | 0.084 | 0.042* |
| Group*Prior | 2.765 | 0.054 | 0.103 |
| Group*Likelihood | 2.829 | 0.056 | 0.099 |
| Group*Prior*Likelihood | 3.405 | 0.066 | 0.071 |

**Table S10:** Between group effects for trial-by-trial sensory weights

| Outcome Variables | Group | *M* | *BF*<sub>01</sub> | *t* | *p* |
| --- | --- | --- | --- | --- | --- |
| P<sub>N</sub>L<sub>N</sub> | NT | 0.056 | 0.798 | 2.026 | 0.048 |
|  | AS | 0.068 |  |  |  |
| P<sub>N</sub>L<sub>W</sub> | NT | 0.042 | 0.966 | 1.910 | 0.062 |
|  | AS | 0.055 |  |  |  |
| P<sub>W</sub>L<sub>N</sub> | NT | 0.021 | 4.222 | 0.510 | 0.613 |
|  | AS | 0.022 |  |  |  |
| P<sub>W</sub>L<sub>W</sub> | NT | 0.040 | 0.380 | 2.429 | 0.019* |
|  | AS | 0.050 |  |  |  |

### 2.0 Continuum Analysis – Autism Traits

We undertook Spearman correlation analysis for each research question.

### 2.1 Main Task

**Table S11:**
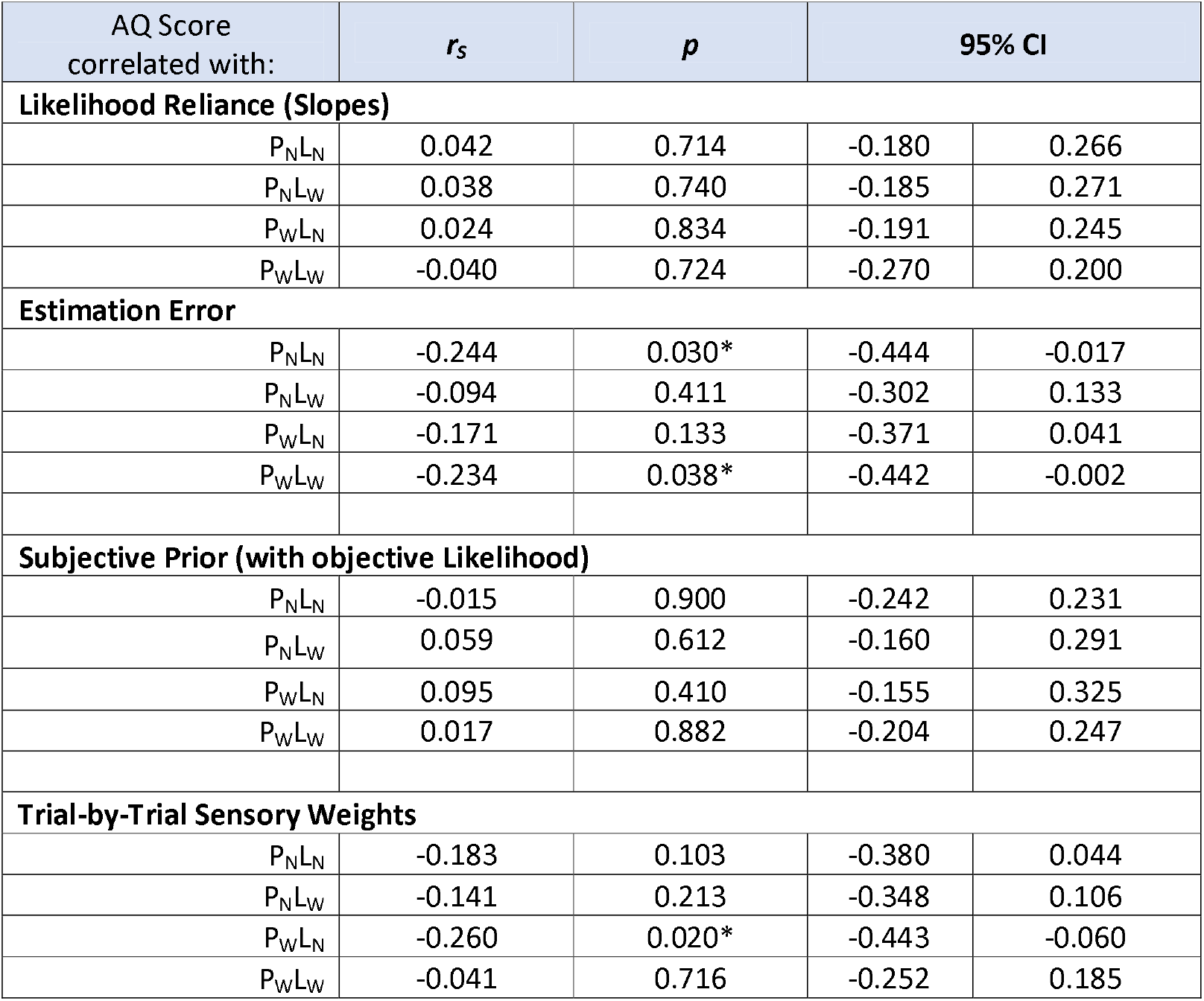
Spearman correlations with Autism Quotient in the main 2-prior task

| AQ Score correlated with: | $r_s$ | $p$ | 95% CI | |
| --- | --- | --- | --- | --- |
| Likelihood Reliance (Slopes) |  |  |  |  |
| $P_{NL_N}$ | 0.042 | 0.714 | -0.180 | 0.266 |
| $P_{NL_W}$ | 0.038 | 0.740 | -0.185 | 0.271 |
| $P_{WL_N}$ | 0.024 | 0.834 | -0.191 | 0.245 |
| $P_{WL_W}$ | -0.040 | 0.724 | -0.270 | 0.200 |
| Estimation Error |  |  |  |  |
| $P_{NL_N}$ | -0.244 | 0.030* | -0.444 | -0.017 |
| $P_{NL_W}$ | -0.094 | 0.411 | -0.302 | 0.133 |
| $P_{WL_N}$ | -0.171 | 0.133 | -0.371 | 0.041 |
| $P_{WL_W}$ | -0.234 | 0.038* | -0.442 | -0.002 |
| Subjective Prior (with objective Likelihood) |  |  |  |  |
| $P_{NL_N}$ | -0.015 | 0.900 | -0.242 | 0.231 |
| $P_{NL_W}$ | 0.059 | 0.612 | -0.160 | 0.291 |
| $P_{WL_N}$ | 0.095 | 0.410 | -0.155 | 0.325 |
| $P_{WL_W}$ | 0.017 | 0.882 | -0.204 | 0.247 |
| Trial-by-Trial Sensory Weights |  |  |  |  |
| $P_{NL_N}$ | -0.183 | 0.103 | -0.380 | 0.044 |
| $P_{NL_W}$ | -0.141 | 0.213 | -0.348 | 0.106 |
| $P_{WL_N}$ | -0.260 | 0.020* | -0.443 | -0.060 |
| $P_{WL_W}$ | -0.041 | 0.716 | -0.252 | 0.185 |

### 2.2 Likelihood Only Task

Spearman rank correlations with AQ scores. For the Likelihood Only Task we excluded outliers based on Tukey’s 1.5 Inter-Quartile Range. Thus 7 participants (1NT and 6AS) were excluded from this analysis.

**Table S12:** Spearman correlations with Autism Quotient in the Likelihood Only Task

|  | NT (N=46) |  |  |  | AS (17) |  |  |  | All (N=73) |  |  |  |
| --- | --- | --- | --- | --- | --- | --- | --- | --- | --- | --- | --- | --- |
| AQ Score correlated with: | r<sub>s</sub> | *p* | 95% CI |  | r<sub>s</sub> | *p* | 95% CI |  | r<sub>s</sub> | *p* | 95% CI |  |
| 1. Accuracy (mean Estimation Error) |  |  |  |  |  |  |  |  |  |  |  |  |
| Narrow | -0.283 | 0.057 | -0.540 | -0.001 | -0.225 | 0.386 | -0.724 | 0.355 | -0.254 | 0.038 | -0.487 | -0.004 |
| Wide | -0.363 | 0.013<br>(0.026)* | -0.621 | -0.094 | -0.158 | 0.560 | -0.665 | 0.412 | -0.359 | 0.003<br>(0.012) * | -0.565 | -0.125 |
| 2. Variance of Estimation Error (i.e., Likelihood Variance) |  |  |  |  |  |  |  |  |  |  |  |  |
| Narrow | -0.051 | 0.736 | -0.307 | 0.229 | 0.004 | 0.987 | -0.575 | 0.557 | 0.062 | 0.618 | -0.175 | 0.289 |
| Wide | -0.303 | 0.041<br>(0.081) | -0.588 | -0.017 | 0.306 | 0.249 | -0.232 | 0.804 | -0.326 | 0.007<br>(0.028) * | -0.542 | -0.082 |

### 3.0 Group by Autism Traits Interaction Effects

We conducted Multivariate analysis of covariance with AQ as covariate, Group as fixed factor and variable interest as the outcome variable(s) to assess the interaction between group and AQ.

### 3.1 Likelihood Only Task

For the Likelihood Only Task we excluded significant outliers based on Tukey’s 1.5 Inter-Quartile Range of sensory weights in the Likelihood Only task. Thus 7 participants (1NT and 6AS) were excluded from the analysis. Thus, the total sample is 73 participants.

**Table S13:** ANCOVA

|  | Outcome variable |  | F | p-value |
| --- | --- | --- | --- | --- |
| 1. | Mean Estimation Error | Narrow Likelihood | 1.757 | 0.165 |
|  |  | Wide Likelihood | 3.503 | 0.020* |
| 2. | Variance of Estimation Error | Narrow Likelihood | 0.435 | 0.728 |
|  |  | Wide Likelihood | 3.107 | 0.033* |

### 3.2 Main / 2-Prior Task

**Table S14:**
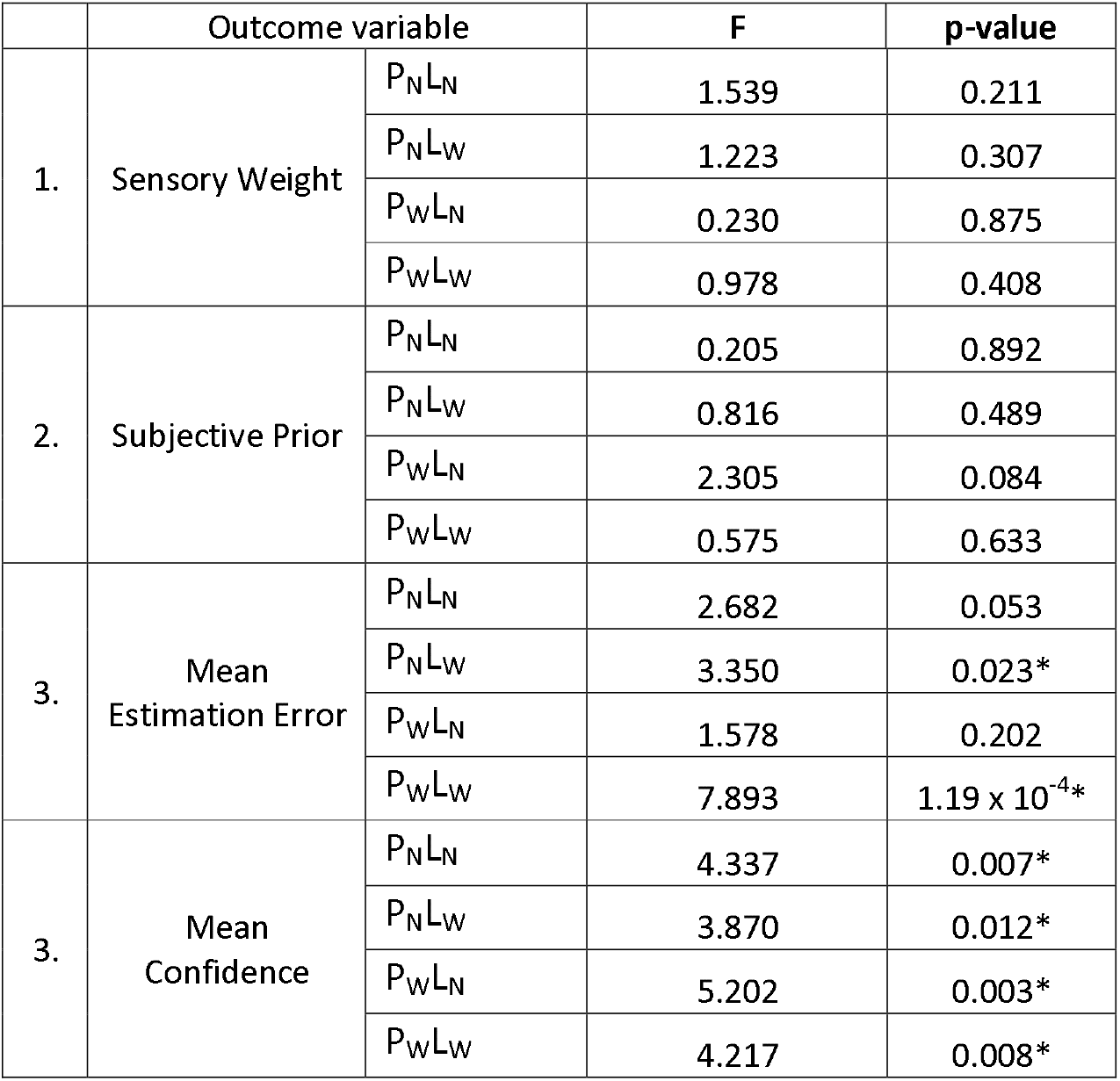
ANCOVA

### 4.0 Autism Subscale Analysis – Attention to detail

**Table S15:** Spearman correlations with AQ subscale attention to detail

| AQ Attention to Detail<br>correlated with: | $r_s$ | p-value | 95% CI | |
| --- | --- | --- | --- | --- |
| Likelihood Only Task: |  |  |  |  |
| 1. Likelihood Variance (RE) |  |  |  |  |
| Narrow | -0.004 | 0.974 | -0.239 | 0.224 |
| Wide | -0.375 | 0.002* | -0.555 | -0.147 |
| 2. Estimation Error |  |  |  |  |
| Narrow | -0.256 | 0.036 | -0.480 | -0.019 |
| Wide | -0.359 | 0.003* | -0.532 | -0.143 |
| 3. Sensory Weight |  |  |  |  |
| Narrow | -0.130 | 0.296 | -0.369 | 0.131 |
| Wide | -0.005 | 0.968 | -0.265 | 0.251 |

### 5.0 Dimensional Analysis – Sensory Sensitivities

Spearman rank correlations with SPQ scores

### 5.1 Main Task

**Table S16:** Spearman correlations with SPQ scores in the 2-Prior Task

| | $r_s$ | $p$ | 95% CI | |
| --- | --- | --- | --- | --- |
| 1. Likelihood Reliance (Slopes) |  |  |  |  |
| $P_{NL_N}$ | -0.091 | 0.425 | -0.311 | 0.154 |
| $P_{NL_W}$ | -0.075 | 0.511 | -0.305 | 0.162 |
| $P_{WL_N}$ | -0.196 | 0.083 | -0.412 | 0.029 |
| $P_{WL_W}$ | -0.109 | 0.338 | -0.321 | 0.126 |
| 2. Estimation Error |  |  |  |  |
| $P_{NL_N}$ | 0.116 | 0.309 | -0.124 | 0.345 |
| $P_{NL_W}$ | 0.017 | 0.885 | -0.227 | 0.250 |
| $P_{WL_N}$ | 0.173 | 0.128 | -0.052 | 0.391 |
| $P_{WL_W}$ | 0.104 | 0.363 | -0.141 | 0.317 |
| 3. Prior Variance |  |  |  |  |
| $P_{NL_N}$ | 0.070 | 0.548 | -0.161 | 0.278 |
| $P_{NL_W}$ | -0.090 | 0.434 | -0.314 | 0.136 |
| $P_{WL_N}$ | -0.217 | 0.058 | -0.420 | 0.005 |
| $P_{WL_W}$ | -0.155 | 0.178 | -0.365 | 0.063 |
| 3. Trial-by-trial sensory weights |  |  |  |  |
| $P_{NL_N}$ | 0.057 | 0.616 | -0.167 | 0.286 |
| $P_{NL_W}$ | 0.009 | 0.937 | -0.199 | 0.235 |
| $P_{WL_N}$ | 0.018 | 0.877 | -0.203 | 0.248 |
| $P_{WL_W}$ | -0.041 | 0.716 | -0.252 | 0.185 |

### 5.2 Likelihood Only Task

**Table S17:**
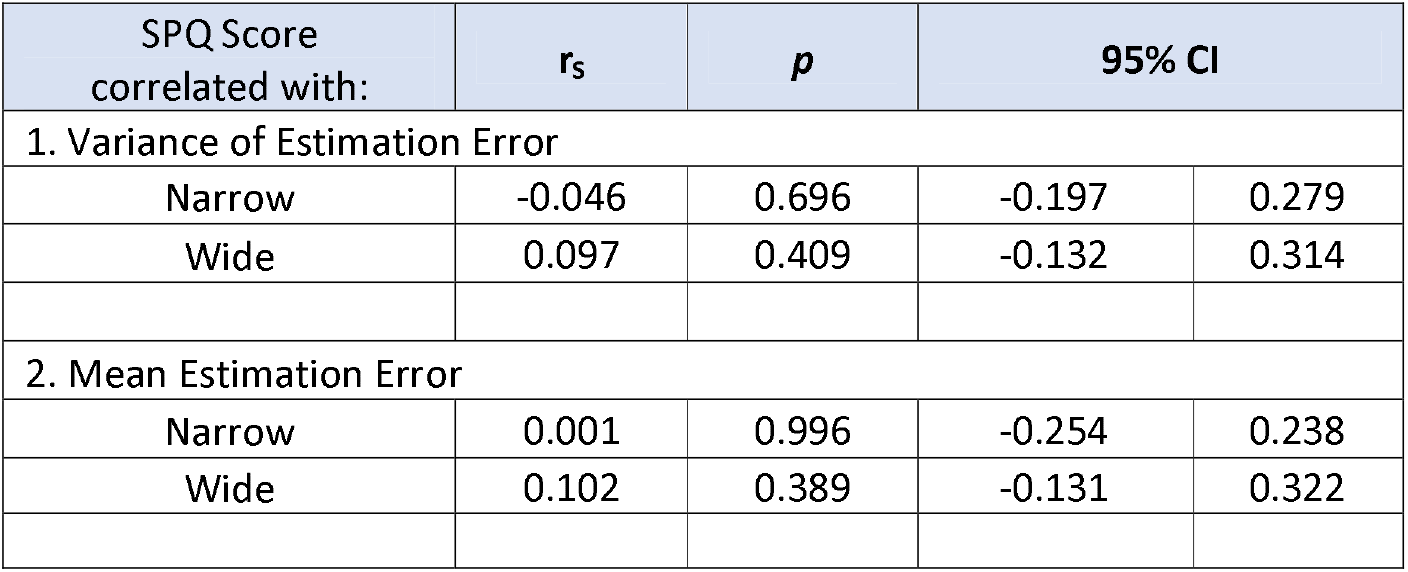
Spearman correlations with SPQ scores in the Likelihood Only Task

| SPQ Score correlated with: | $r_s$ | $p$ | 95% CI | |
| --- | --- | --- | --- | --- |
| 1. Variance of Estimation Error |  |  |  |  |
| Narrow | -0.046 | 0.696 | -0.197 | 0.279 |
| Wide | 0.097 | 0.409 | -0.132 | 0.314 |
| 2. Mean Estimation Error |  |  |  |  |
| Narrow | 0.001 | 0.996 | -0.254 | 0.238 |
| Wide | 0.102 | 0.389 | -0.131 | 0.322 |

### 6.0 ADOS Scores

Bootstrapped Spearman rank correlations with ADOS scores

**Table S18:** ADOS Scores and behavioural measures

| ADOS Score correlated with: | $r_s$ | p-value | 95% CI | |
| --- | --- | --- | --- | --- |
| Likelihood Only Task: |  |  |  |  |
| 4. Likelihood Variance (RE) |  |  |  |  |
| Narrow | 0.080 | 0.687 | -0.326 | 0.447 |
| Wide | 0.099 | 0.616 | -0.336 | 0.489 |
| 5. Estimation Error |  |  |  |  |
| Narrow | 0.186 | 0.353 | -0.247 | 0.581 |
| Wide | 0.105 | 0.602 | -0.329 | 0.529 |
| 6. Likelihood Reliance |  |  |  |  |
| Narrow | 0.098 | 0.626 | -0.363 | 0.563 |
| Wide | -0.164 | 0.413 | -0.516 | 0.263 |
| 2-Prior Task: |  |  |  |  |
| 1. Subjective prior variance |  |  |  |  |
| $P_N L_N$ | 0.079 | 0.674 | -0.267 | 0.405 |
| $P_N L_W$ | -0.014 | 0.939 | -0.375 | 0.333 |
| $P_W L_N$ | -0.277 | 0.131 | -0.626 | 0.143 |
| $P_W L_W$ | -0.266 | 0.148 | -0.567 | 0.135 |
| 2. Estimation Error |  |  |  |  |
| $P_N L_N$ | 0.256 | 0.164 | -0.138 | 0.621 |
| $P_N L_W$ | 0.186 | 0.317 | -0.178 | 0.524 |
| $P_{WL_N}$ | 0.199 | 0.282 | -0.214 | 0.545 |
| $P_{WL_W}$ | 0.284 | 0.122 | -0.110 | 0.630 |

### 7.0 Regression model fit tests

We investigated if the regression model fit was significant for each condition and each participant. We find the regression model was significant (p<0.5) for each condition for most participants.

However, 9 (out of 80) participants showed non-significant p-values (p<0.05) in one or two conditions but were significant for the other conditions. (These 9 participants included 5 ‘NT’, 4 ‘AS’ and 2 ‘Other’). We have not excluded these for any analysis.

2 subjects in the AS group (i.e., with a confirmed diagnosis of ASD) however, showed non-significant p-values for all four conditions. These participant’s parameter estimates are not outliers amongst other participants. However, we have conducted the group analysis that impacts our major conclusions excluding these two participants and found similar results, hence we decided to keep the original results with all participants included in the main text. Below are the results excluding the two participants from group analysis:

1) Likelihood reliance in the 2-prior task

We found evidence for no difference between the groups in overall likelihood reliance, (i.e., sensory weights averaged across conditions) [*t* = -0.448, BF_01_ = 4.256, *p* = 0.656].
2) Context adjustment in the 2-prior task

Looking within conditions in the 2-Prior Task, a repeated-measures ANOVA (see methods) of sensory weights revealed a significant Group*Prior [Effect Size η_p_^2^ = 0.109, p = 0.022], a non-significant trend in Group*Likelihood [η_p_^2^ = 0.067, p = 0.071] interaction, but no Group*Prior*Likelihood [η_p_^2^ = 0.027, p = 0.260] interaction.

